# Amalga: Designable Protein Backbone Generation with Folding and Inverse Folding Guidance

**DOI:** 10.1101/2023.11.07.565939

**Authors:** Shugao Chen, Ziyao Li, Xiangxiang Zeng, Guolin Ke

**Author notes:** Corresponding author: Ziyao Li < >. This authors contributed equally to this paper. Shugao Chen contributed as an intern at DP Technology.

## Abstract

Recent advances in deep learning enable new approaches to protein design through inverse folding and backbone generation. However, backbone generators may produce structures that inverse folding struggles to identify sequences for, indicating designability issues. We propose Amalga, an inference-time technique that enhances designability of backbone generators. Amalga leverages folding and inverse folding models to guide backbone generation towards more designable conformations by incorporating “folded-from-inverse-folded” (FIF) structures. To generate FIF structures, possible sequences are predicted from step-wise predictions in the reverse diffusion and further folded into new backbones. Being intrinsically designable, the FIF structures guide the generated backbones to a more designable distribution. Experiments on both de novo design and motif-scaffolding demonstrate improved designability and diversity with Amalga on RFdiffusion.

## 1 Introduction

Rational protein design aims to create novel proteins or modify existing ones to obtain desired structures and functions. Accurate protein design methods enable direct applications such as enzyme engineering [10] and antibody-based drug design [15]. However, the vast combinatorial spaces of protein sequence and structure, along with their intricate interdependence, render this problem a longstanding challenge in biotechnology.

Fortunately, recent advances in deep learning illuminate new approaches to design proteins *de novo*. Capitalizing on abundant sequence and structure data, *inverse protein folding models* [7, 4] have succeeded in designing protein sequences that fold into specified target structures. Meanwhile, inspired by the formidable successes of diffusion models in image generation [6, 11], *diffusion-based backbone generators* [2, 13, 16, 17, 14] explore the prospects of generating novel protein backbone structures. The integration of these two methods outlines a pipeline to design proteins: 1) sample protein structures using backbone generators; 2) determine corresponding sequences with inverse folding models; 3) screen the generated proteins based on *designability* - how well the generated sequence folds into the accompanying structure; and 4) further screen the *designable* structures for desired applications, based on both sequences and structures.

While existing backbone generation models, as exemplified by RFdiffusion [14], produce backbones with sensible local structures and appropriate proportions of stable secondary structures (helices and sheets), inverse folding models struggle to identify sequences for a sizable proportion of the generated backbones, even when human evaluation deeming them reasonable. Quantitatively, RFdiffusion benchmarking indicates approximately 30% of samples did not satisfy the designability criterion. We reckon this issue arises due to two possible factors: 1) current protein folding and inverse folding models lack sufficient accuracy; 2) most existing backbone generation models are explicitly trained to reproduce structures alone, without capturing the intricate sequence-structure relationship which essentially depicts designability. We aim to address the second factor in this paper.

Here we propose Amalga, a simple yet effective inference-time technique to enhance the designability of diffusion-based backbone generators. By harnessing off-the-shelf folding and inverse folding models, Amalga guides backbone generation towards more designable conformations. Specifically, Amalga generates a set of “folded-from-inverse-folded” (FIF) structures by folding the sequences which are inverse folded from step-wise predicted backbones. These FIF structures, being inherently designable, are aligned to the predicted backbone and input into RFdiffusion’s self-conditioning channel. Intuitively, this encourages RFdiffusion to match the distribution of designable structures. While retraining or finetuning RFdiffusion with FIF inputs may further improve performance, we demonstrate that Amalga significantly boosts designability when applied solely during inference.

## 2 Preliminaries

### Diffusion-based Protein Backbone Generation

Recent works [1, 13, 17] have explored generating protein backbones using diffusion models such as denoising diffusion probabilistic model (DDPM) [6] and generative stochastic differential equations [12]. These generative models leverage forward and reverse diffusion processes to gradually transform samples from a simple prior distribution (often Gaussian) into complex backbone structures. The forward process perturbs the coordinates and orientations of each residue by adding noise with different scales on the timestep *t*. The reverse process then recovers high-quality backbones by iteratively predicting less noisy versions from the prior. Taking DDPM as an example, the forward and backward diffusion processes are formulated as:

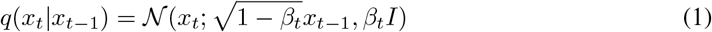

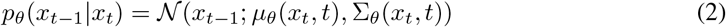

where *q*(*x*_*t*_ |*x*_*t* −1_) perturbs *x*_*t* −1_ with Gaussian noise to obtain *x*_*t*_, and *p*_*θ*_ predicts the reverse step result with neural networks based on the diffused sample *x*_*t*_.

### Rfdiffusion

RFdiffusion [14] is a recent example of diffusion-based backbone generators. It finetunes RosseTTAFold [3], a multiple sequence alignment (MSA) based protein structure prediction model, with noised samples generated from the forward diffusion. Specifically, the structures of proteins to be designed are perturbed, and their corresponding sequences are masked. RFdiffusion also utilizes the original template channel in RosseTTAFold to input previously generated backbones into the model for self-conditioning. In this work, we additionally send the generated FIF samples through this channel to encourage the model to match the distribution of designable structures.

### Designability Formulation

Given a folding model *f* and an inverse folding model *f* ^−1^, the general designability metric D(**x**) is defined as:

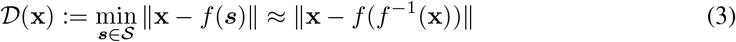

where *s* and x denote protein sequences and structures respectively, ∥. ∥quantifies the structural differences between two conformations (e.g. RMSD), and S represents the set of all feasible protein sequences. Conceptually, designability measures how accurately a structure x can be reproduced by its predicted sequence *f* ^−1^(x) after folding, with lower values indicating higher designability. Notably, this metric depends on the accuracy of the folding and inverse folding models.

## 3 Method

Figure 1 illustrates the workflow of Amalga at each timestep *t* of the reverse diffusion process. We demonstrate Amalga on RFdiffusion, while the idea is broadly applicable to other baselines [17, 13]. The model takes as input the noised backbone x_*t*+1_ from the forward process, the predicted backbone 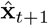, and the FIF samples 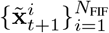 generated in the previous step to make new backbone 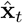. This predicted backbone, together with x_*t*+1_, is used to compute the noised input for the next timestep via the reverse diffusion formula. Amalga then inverse folds the prediction 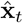 using ESM-IF [7] to generate possible sequences 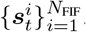. These sequences are then folded using ESMFold [9] to obtain new FIF backbones 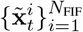 to guide the next step. Note that 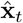 is directly produced by the backbone generator, thus assumed to have more reasonable structures and sensible inverse folding results, while x_*t*_ is the intermediate result in the diffusion process, thus with discontinuities.

**Figure 1.**
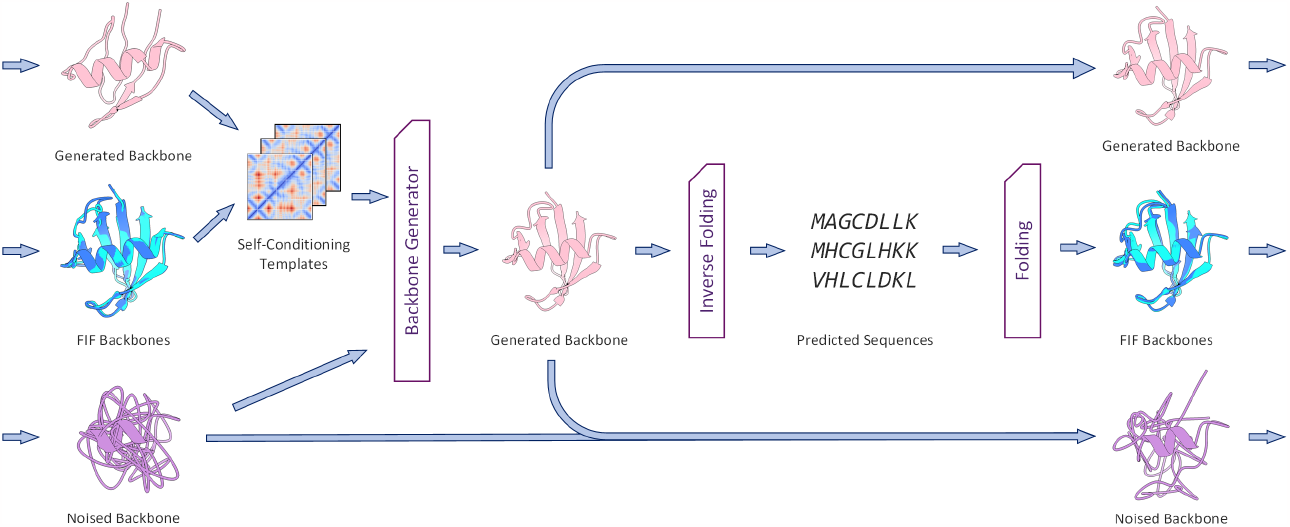
Amalga pipeline in each step. Amalga generates FIF samples from step-wise predicted backbones and inputs them to the model via the self-conditioning channel.

In an attempt to further guide the model towards predicting foldable protein structures, we also explored providing the predicted sequences 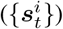 as inputs to the sequence channel of RFdiffusion. As the sequence channel in the original RosseTTAFold architecture leveraged multiple sequence alignments (MSAs) to inform structural predictions, we hypothesized that explicitly providing these inverse-folded sequences could similarly enhance folding precision. However, as is shown in the next section, experiments inserting 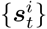 did not always result in improved performance. We believe the model likely requires a finetuning stage in order to integrate the sequences more properly.

In fact, the FIF samples directly inherit zero designability error, since their sequences are known to fold into the generated structures (up to the error of the folding model). As such, they are already successful design outcomes for unconstrained, *de novo* generation. However, these FIF structures may not conform to the motif constraints specified by the user. In contrast, the final output structure from the reverse diffusion process will explicitly satisfy the desired motifs, since they are fixed during this process. Therefore, Amalga balances global designability, provided by guiding the diffusion model with the FIF samples, and precise motif reconstruction in the final output.

## 4 Experiments

### Settings

We conducted comparative experiments between the original RFdiffusion model and the RFdiffusion model augmented with Amalga. We utilized ProteinMPNN [4] following RFdiffusion for inverse folding, however, we replaced AlphaFold [8] with ESMFold to fold the final structures, as ESMFold achieves superior performance when multiple sequence alignments are unavailable. This replacement did not significantly altered the results, as we have analyzed in the appendix. For Amalga, we tested settings with *N*_FIF_ = 1, 5. We reported results for two Amalga variants: one where we input predicted sequences via the MSA channel (denoted “+seq”), and one where we did not.

We evaluated two backbone generation task schemes: 1) de novo design, in which backbones are generated without external constraints, and 2) motif-scaffolding, in which backbones should contain a predefined motif with known sequence and structure. In the former task, we generated 20 structures of lengths 100, 150, 200, 250 and 300, respectively. In the latter task, we generated 100 structures for each of the 25 benchmark tasks in RFdiffusion. We use the root-mean-square deviation of self-consistency (scRMSD) and the in silico success rate to depict designability. The scRMSD measures the error between a generated structure and its closest foldable structure:

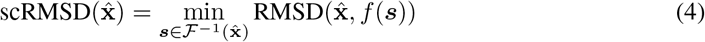

where 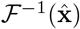 denotes the set of 8 inverse-folded sequences of 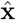. The in silico success criteria were adopted from RFdiffusion: for de novo design, an scRMSD below 2Å was required to be considered successful; for motif-scaffolding, an additional requirement was that the RMSD between the motif in the best design and the target motif be less than 1Å. We also report a metric of design diversity: all success backbones were clustered using MaxCluster [5] with a TM-score threshold of 0.5. The diversity was quantified as the number of the unique clusters in the generated success samples.

### Results

Table 1 shows metrics averaged on all 100 de novo generation samples and 2500 motif-scaffolding samples. Notably, results for motif-scaffolding are more informative, as in de novo task the FIF samples are already samples with zero scRMSD. Overall, Amalga obtained superior designability and diversity over RFdiffusion. Adding the sequence into the MSA channel (+seq) with *N*_FIF_ = 1 improves the performance, while the contradictory result holds with *N*_FIF_ = 5. We posit that the model needs further training to adapt to more inverse-folded sequences. We examine the specific success rate of 25 motif-scaffolding tasks in Figure 2. Amalga consistently outperforms RFdiffusion on the 25 benchmark cases with few exceptions. Notably, we observed significant improvements in the RFdiffusion performance over the originally reported. We posit the current release of RFdiffusion parameters have been refined since its publication. Results with regard to motif RMSDs, etc. are available in the appendix.

**Table 1:** Results of Amalga and the original RFdiffusion, averaged over all cases.

|  |  | DE NOVO |  |  | MOTIF-SCAFFOLDING |  |  |
| --- | --- | --- | --- | --- | --- | --- | --- |
|  |  | % success | scRMSD | Diversity | % success | scRMSD | Diversity |
| RFdiffusion |  | 86.00 | **1.29** | 15.60 | 69.28 | 1.24 | 14.24 |
| Amalga | ($N_{\text{FIF}} = 1$) | 86.00 | 1.58 | **16.00** | 73.20 | 1.05 | 14.88 |
| | ($N_{\text{FIF}} = 1$, +seq) | **87.00** | 1.30 | 15.40 | 74.24 | **0.98** | 14.24 |
| | ($N_{\text{FIF}} = 5$) | 83.00 | 1.65 | 15.40 | **75.64** | **0.98** | **16.56** |
| | ($N_{\text{FIF}} = 5$, +seq) | 83.00 | 1.43 | 15.80 | 70.84 | 1.03 | 16.44 |

**Table 2:**
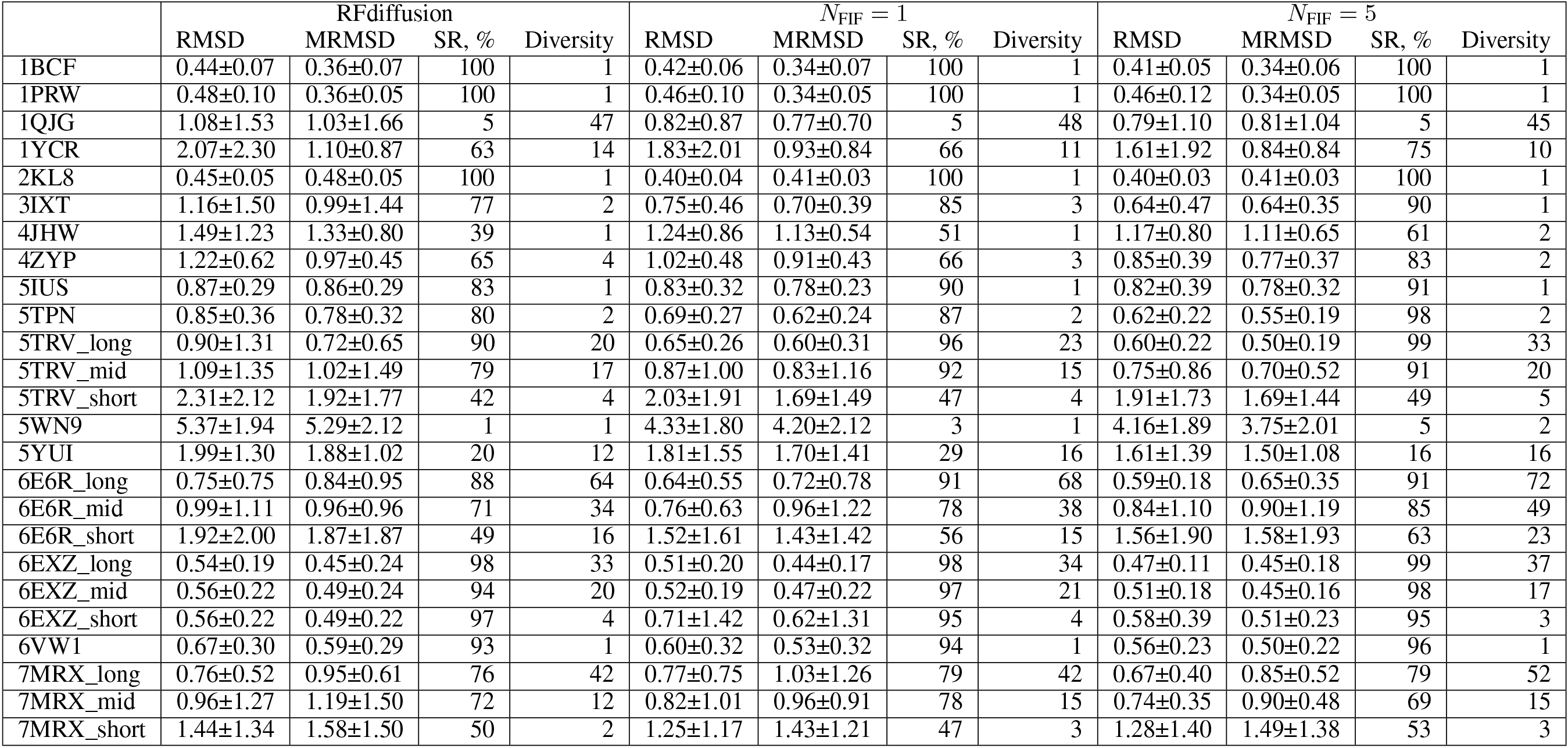
Detailed results of 25 benchmark tasks without additional sequences. MRMSD refers to the motif RMSD and SR refers to the success rate.

**Table 3:**
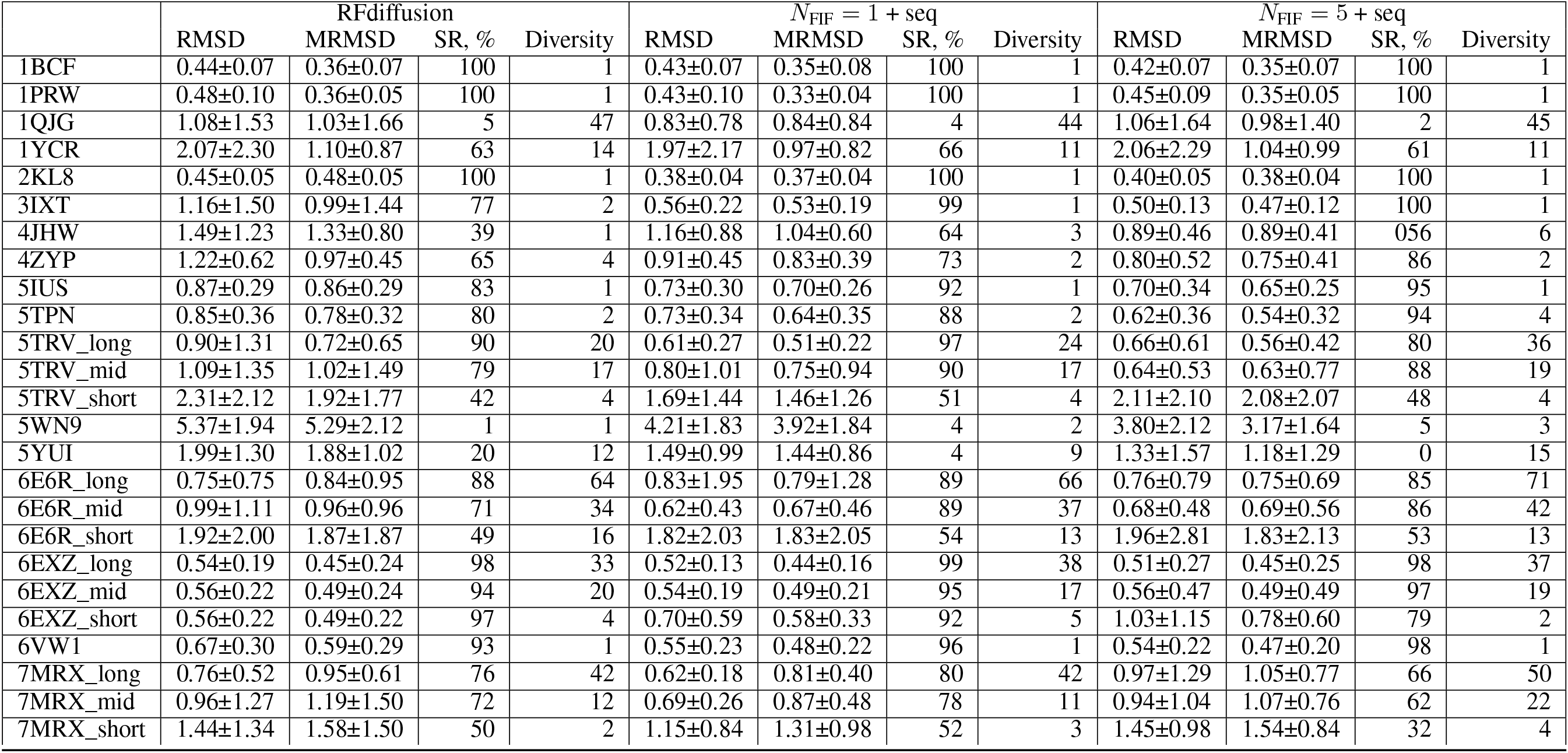
Detailed results of 25 benchmark tasks with additional sequences. MRMSD refers to the motif RMSD and SR refers to the success rate.

**Figure 2.**
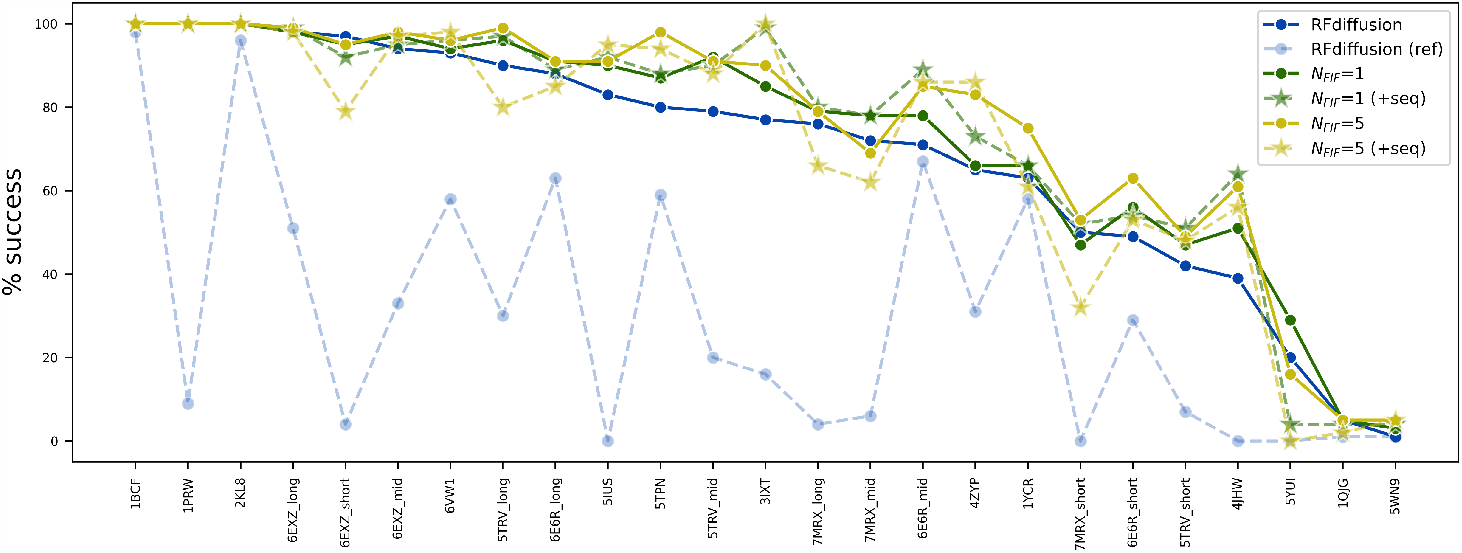
Success rate of 25 benchmark tasks in RFdiffusion. RFdiffusion (ref) in light blue displays statistics directly taken from [14], while RFdiffusion in dark blue shows reproduced statistics under our settings, comparable to Amalga. The columns are ranked by RFdiffusion performance.

### Efficiency

We analyzed the running time of Amalga to quantify the introduced complexity of FIF computation. To generate one 100 amino acid protein, the running time on a 32GB NVIDIA V100 GPU increased from 1^′^00^′′^ to 3^′^04^′′^ with *N*_FIF_ = 1 and 5^′^12^′′^ with *N*_FIF_ = 5. Overall, the introduced complexity is comparable to the original model’s complexity.

## 5 Conclusion & Future Work

In this work, we have proposed Amalga as a broadly applicable inference-time technique to enhance the designability of diffusion-based backbone generators exemplified by RFdiffusion. Our experiments demonstrate that Amalga successfully improves the designability and diversity of generated structures from RFdiffusion, at the cost of additional inverse folding and folding computations. As a direct path for improvement, an obvious next step is to fine-tune RFdiffusion to better adapt it to Amalga inputs. Furthermore, inference speed could be enhanced by optimizing Amalga implementation, such as enabling batched ESMFold inference. Since predicted backbones may not vary drastically step-by-step, utilizing longer intervals between FIF evaluations leads to another gain of efficiency. For more rigorous validation, pending experimental conditions, we hope to perform wet lab experiments to further prove the effectiveness of Amalga designs. As ongoing work, we are also actively exploring adaptations of this approach to other existing protein design baselines.

## A Implementation Details

Our work implements the open-source code of RFdiffusion ^3^. We use the default sampling settings from RFdiffusion, except where noted. The number of sampling steps is set to 50, and the *C*_*α*_ translation noise scalar is 1. For generating FIF samples, we leverage ESM-IF and ESMFold models ^4^ due to their state-of-the-art performance and efficiency. To evaluate generated sequences orthogonally, we follow the original RFdiffusion and use ProteinMPNN ^5^, replacing the protein folding model from AlphaFold with ESMFold for its superior single-sequence structure prediction.

## B Effect of Folding Models

To examine whether the folding model used for evaluation impacts the final results, we generate 10 samples for each case and fold the same ProteinMPNN sequences with both ESMFold and Alphafold2. Using the same designability criteria as described in Section 4, Fig. 3 shows that the choice of folding model does not substantially influence the evaluation of designability.

**Figure 3.**
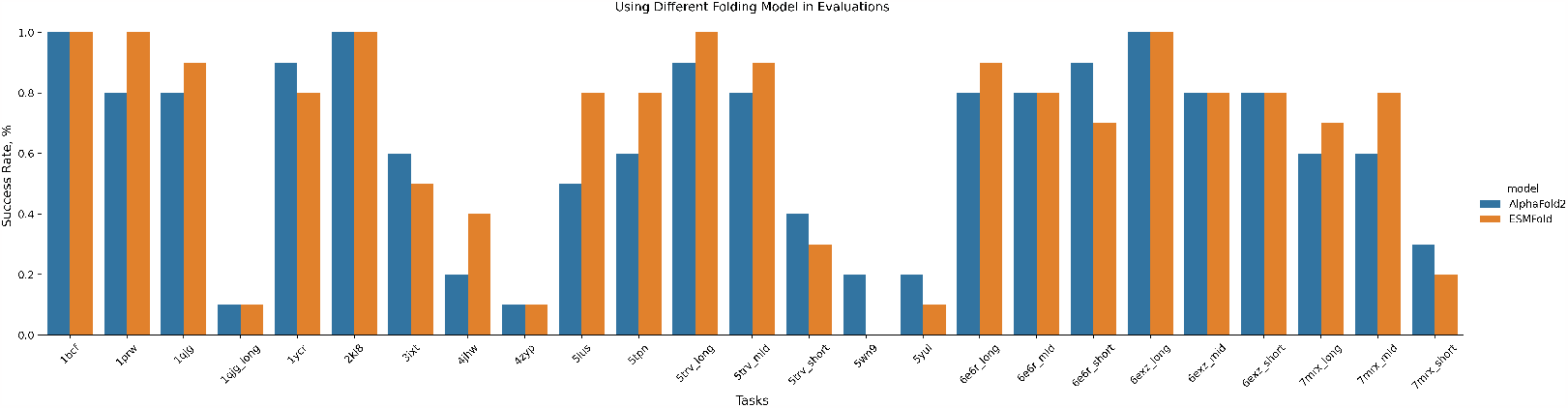
Designability metrics between AlphaFold and ESMFold.

## C RMSD Variation in Sampling

We plot the step-wise RMSD and motif RMSD between the backbone generator output 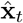 and the FIF samples 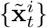 during the denoising process on two motif-scaffolding cases. Notably, as 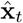 maintains the target motif almost identically, the reported motif RMSD reflects the deviation between FIF samples and the target motif. As shown in Figure 4, both RMSDs decrease consistently following the reverse process.

**Figure 4.**
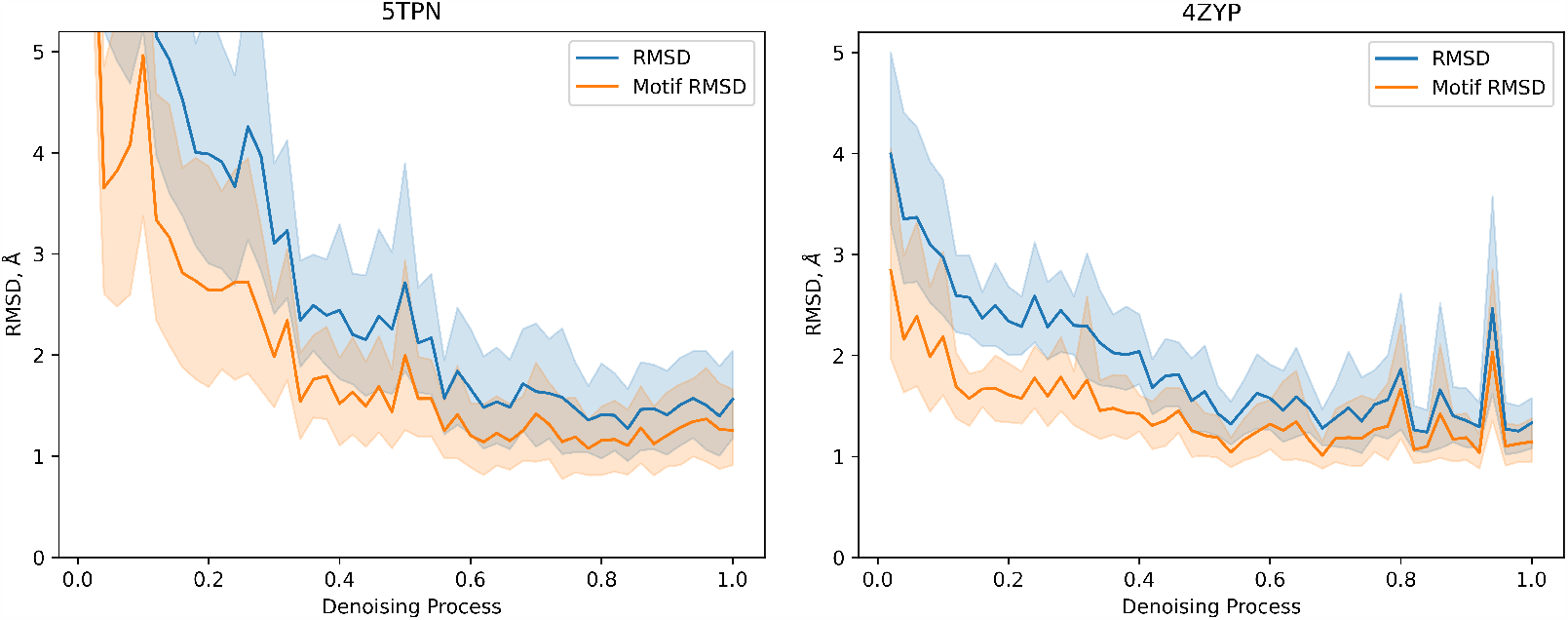
Step-wise RMSDs between FIF samples and generated backbones.

## Footnotes

3 https://github.com/RosettaCommons/RFdiffusion

4 https://github.com/facebookresearch/esm

5 https://github.com/dauparas/ProteinMPNN

